# Expanding gmx_MMPBSA capabilities for end-state binding free energy calculations with GROMACS and AMBER

**DOI:** 10.64898/2026.10.24.761505

**Authors:** Mario E. Valdés-Tresanco, Mario S. Valdés-Tresanco, Glenda Romero-Hernández, Pedro A. Valiente, Ernesto Moreno

## Abstract

gmx_MMPBSA is an end-to-end tool for performing end-state binding free energy calculations. Here, we present advances in the gmx_MMPBSA suite that simplify preparation, expand the calculation options, and improve the analysis of results. The new features include the GBNSR6 and nonlinear PB models, second-order cumulant (C2) entropy estimation, inclusion of explicit structural waters, and multiple alanine/glycine mutations. We also introduce amber_MMPBSA, a new entry point that accepts native AMBER input files and extends the calculation and analysis capabilities of gmx_MMPBSA to AMBER users. The redesigned graphical analyzer, gmx_MMPBSA_ana, and modernized Python API facilitate exploration of large datasets and comparisons across multiple systems. We illustrate the new gmx_MMPBSA capabilities using a diverse set of biomolecular complexes (protein-protein, protein-RNA, protein-small molecule, DNA-small molecule) and show how the calculated binding energies vary across systems depending on the methodological choices. The source code for the latest version 1.7.1 is available at https://github.com/Valdes-Tresanco-MS/gmx_MMPBSA.

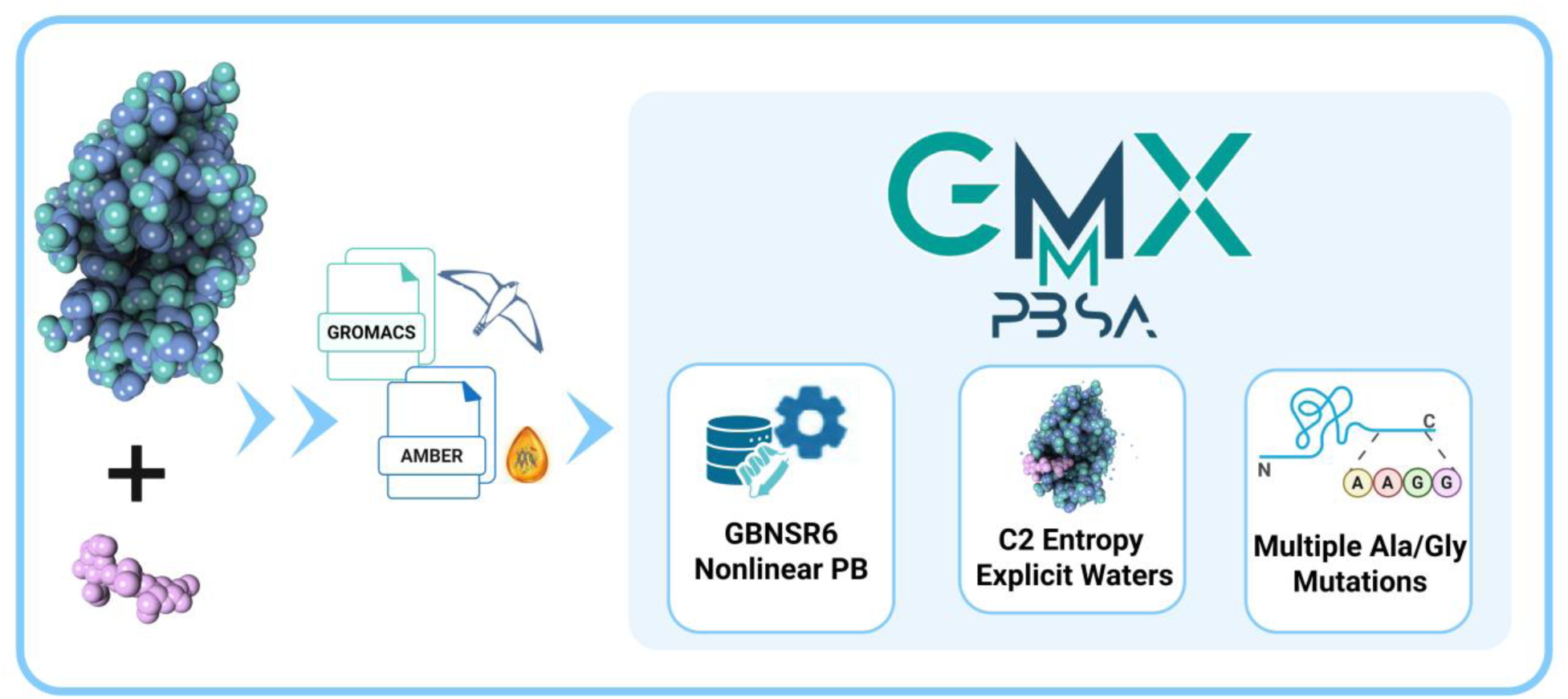

## Introduction

Molecular Mechanics/Poisson-Boltzmann (Generalized Born) Surface Area (MM/PB(GB)SA) methods estimate binding free energies from molecular ensembles by combining molecular-mechanics energies, polar and nonpolar solvation contributions, and entropy estimates.^1–4^ Their relatively low computational cost and applicability to diverse systems have made them widely used to study protein-ligand,^5–7^ protein-protein,^8–10^ and protein-nucleic acid^11,12^ complexes. We originally developed the gmx_MMPBSA suite to perform these calculations using GROMACS trajectories. The gmx_MMPBSA suite included an installation-testing tool, gmx_MMPBSA_test, and a dedicated graphical analyzer, gmx_MMPBSA_ana.^13^

To date, the gmx_MMPBSA suite has been widely adopted within the GROMACS community, with more than 3,400 citations as of October 2026. However, earlier releases presented practical limitations in input compatibility, calculation setup, and analysis. For instance, former gmx_MMPBSA versions were intended for working specifically with GROMACS files,^14^ which severely restricted its application to systems run with other simulation engines *(e.g.*, AMBER, NAMD, OpenMM). The initial versions of the calculation engine limited access to most parameters controlling PB and 3D-Reference Interaction Site Model (3D-RISM) calculations and lacked support for newer solvation models *(i.e.*, GBNSR6^15^) and more sophisticated methods such as second-order cumulant (C2) entropy estimation, inclusion of explicit structural waters, or multiple alanine/glycine scanning. Furthermore, membrane placement and definition of the complex, receptor, and ligand QM-region charges in QM/MMGBSA calculations relied on manual assignment, increasing the risk of setup errors. Finally, inefficiencies in the initial version of the analyzer gmx_MMPBSA_ana complicated the analysis of large datasets and comparisons across systems.

Here, we present extensions to the gmx_MMPBSA suite that address these limitations. We demonstrate the expanded gmx_MMPBSA workflow using representative biomolecular complexes and examine how methodological choices affect the resulting binding energy estimates.

## Improvements to the gmx_MMPBSA workflow

### The gmx_MMPBSA workflow

The gmx_MMPBSA suite provides an end-to-end workflow to perform end-state binding free energy calculations through three main steps: *i*) automated topology preparation, *ii*) energy evaluation using implicit solvent models, and *iii*) visualization and analysis of the results (Figure 1).

**Figure 1.**
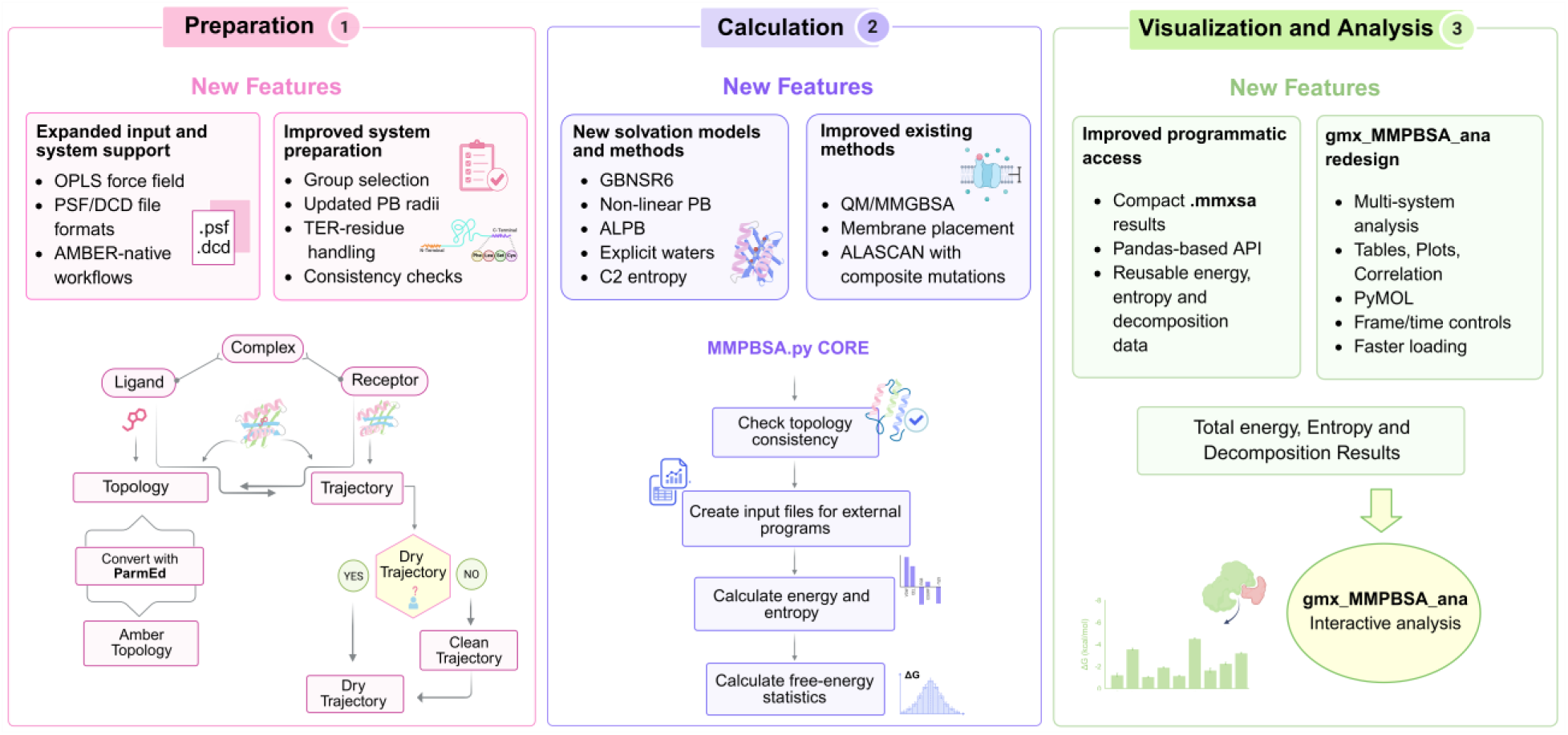
gmx_MMPBSA workflow. The workflow comprises three stages: preparation of receptor-ligand inputs, end-state free energy calculation, and visualization and analysis. The lower portions of the panels show the initial workflow; the upper portions summarize the new features, including broader input support, improved system preparation, new solvation models and methods, and a redesigned graphical analyzer (gmx_MMPBSA_ana) and improved programmatic access to results. Created in BioRender. Valdes, M. (2026) https://BioRender.com/rln2tpv

Briefly, in the preparation stage, gmx_MMPBSA reads the molecular structure, topology, and molecular dynamics (MD) trajectory; validates the receptor and ligand selections; converts the GROMACS topology into an AMBER-compatible representation through ParmEd; and extracts the selected trajectory frames. The prepared inputs are then used for end-state energy calculations with GB, PB, or 3D-RISM solvent models. The resulting data can be explored interactively using the graphical analyzer gmx_MMPBSA_ana or accessed programmatically through the Python API.

### Topology preparation and input support

The initial version of gmx_MMPBSA supported AMBER- and CHARMM-derived GROMACS topologies. Subsequent releases extended this support to OPLS-derived topologies and introduced a documented workflow for converting PSF/DCD files from systems prepared for NAMD, OpenMM, and GENESIS into the GROMACS inputs required by gmx_MMPBSA. Earlier versions also provided a structure-based route using AMBER’s tleap to generate AMBER-compatible topologies. However, this approach had limitations for complex systems, particularly those containing components that required parameterization with third-party software. Beginning with version 1.7.0, gmx_MMPBSA requires the original GROMACS topology and associated parameter files, so that topology preparation uses the same force-field parameters supplied for the MD simulation.

Version 1.7.0 also introduced amber_MMPBSA (https://valdes-tresanco-ms.github.io/gmx_MMPBSA/1.7.0/amber_MMPBSA), a preparation route that accepts native AMBER input files (Figures 1 and 2). amber_MMPBSA reads AMBER topology and trajectory files directly and uses AMBER masks to select the receptor and ligand, bypassing conversion through GROMACS. This route preserves the implicit-solvent atomic radii and screening factors that are typically stored in the AMBER topologies. After preparation, amber_MMPBSA uses the same calculation, output, graphical analysis, and Python API framework as gmx_MMPBSA.

**Figure 2.**
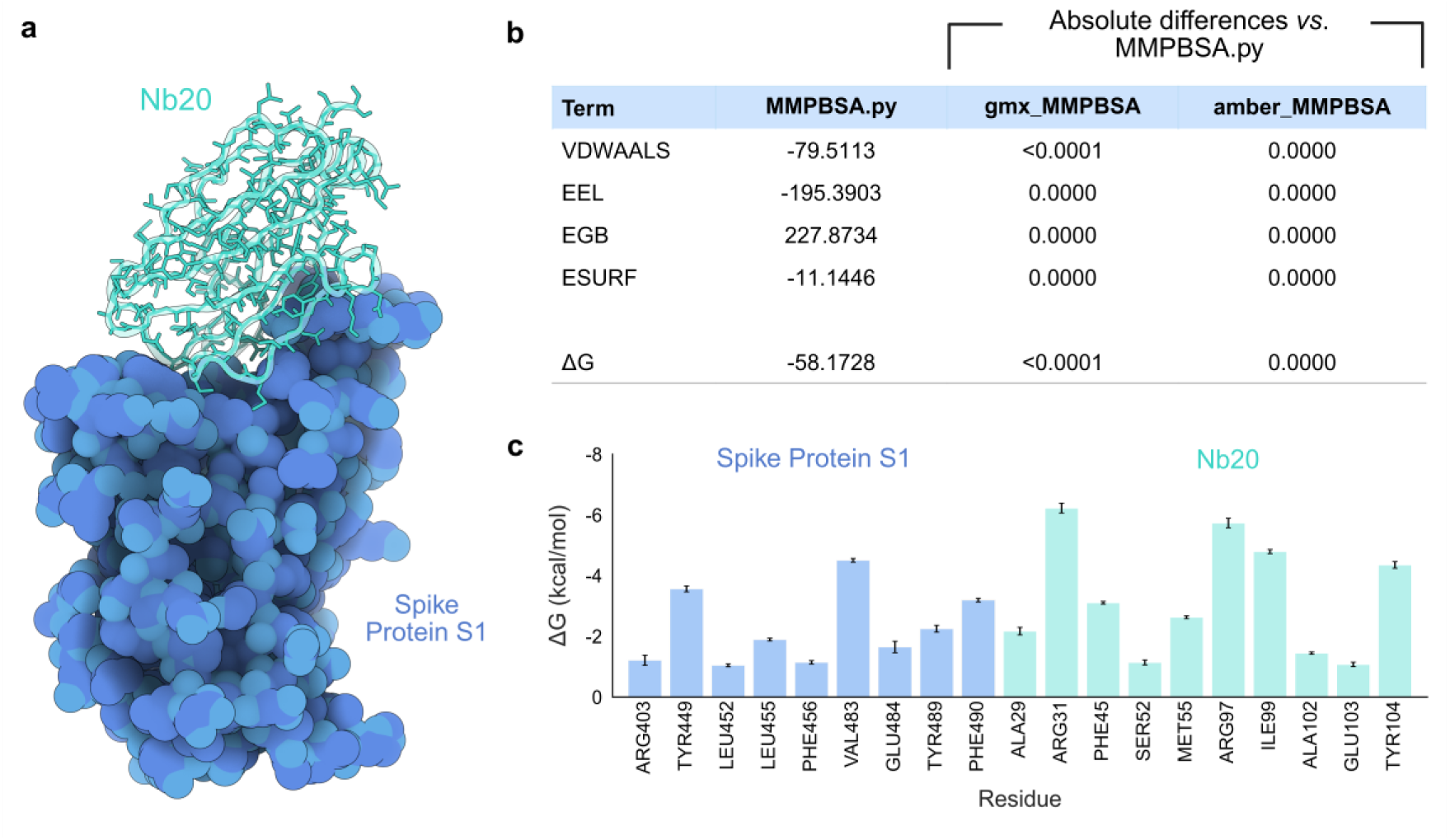
Comparison of MM/GBSA energies calculated with gmx_MMPBSA, amber_MMPBSA, and MMPBSA.py. **a.** Three-dimensional representation of the nanobody Nb20 bound to the spike protein S1 from the SARS-CoV-2 virus (PDB ID: 7JVB). **b.** Absolute differences (in kcal/mol) in nonbonded binding-energy terms calculated with the GB-Neck2 model (igb=8) in gmx_MMPBSA and amber_MMPBSA relative to MMPBSA.py. Bonded terms are omitted because they cancel when using the single-trajectory protocol. **c.** Per-residue energy decomposition for interfacial residues calculated with amber_MMPBSA. Error bars represent standard errors. Created in BioRender. Valdes, M. (2026) https://BioRender.com/1yz2spr

To verify the numerical consistency of the new amber_MMPBSA implementation, we compared MM/GBSA energies calculated with amber_MMPBSA, gmx_MMPBSA, and MMPBSA.py for the SARS-CoV-2 spike receptor-binding domain complexed with nanobody Nb20 (PDB ID: 7JVB).^16^ amber_MMPBSA reproduced the energies obtained with gmx_MMPBSA and MMPBSA.py (Figure 2). Furthermore, we performed per-residue decomposition and observed substantial contributions to the binding energy from previously described interfacial residues.^16^ For example, residues Glu484 and Phe490 from the spike protein, together with Ala29, Arg31, Met55, Arg97, and Tyr104 from the Nb20 nanobody, were involved in a polar interaction network and showed substantial contributions (> 1 kcal/mol in magnitude) to the binding energy. Residues Leu452, Val483, and Phe490 from the spike protein, together with Phe45 and Met55 from Nb20, belong to the hydrophobic interaction network^16^ and also showed substantial binding-energy contributions (Figure 2).

### Calculation methods

Recent developments in gmx_MMPBSA (also available in amber_MMPBSA) have expanded end-state binding free energy calculations to include new solvation models (*e.g.*, GBNSR6) and methods (*e.g.*, ALPB correction, explicit waters). These methods can be combined with existing, well-established solvation models such as GB-OBC2, as well as with the newly implemented solvation models (Figure 3). In addition, several existing methods, including QM/MMGBSA and membrane calculations, have been improved to simplify the calculation setup and reduce the risk of setup errors.

**Figure 3.**
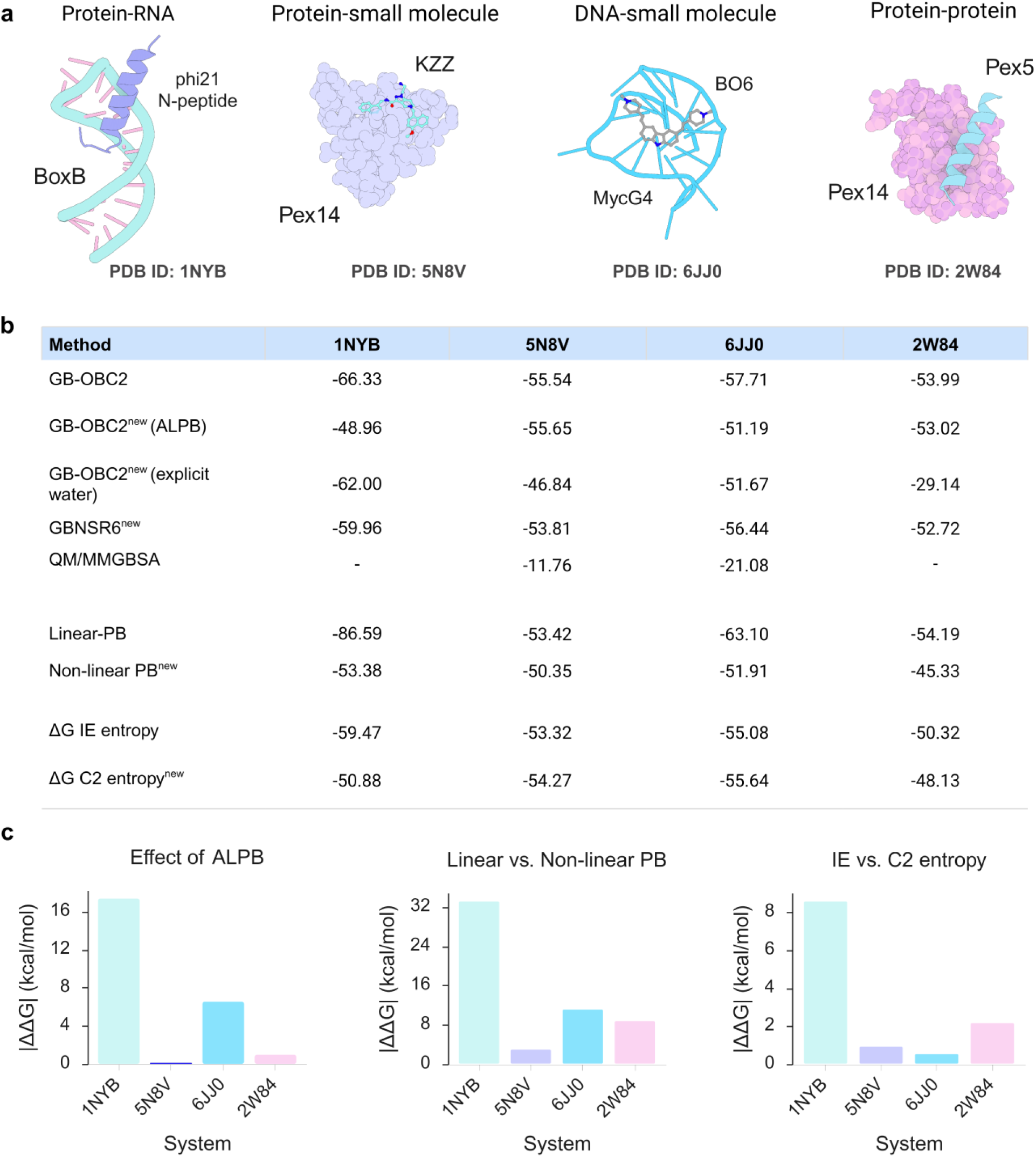
Application of established and newly implemented calculation options in gmx_MMPBSA to four biomolecular complexes. **a.** Three-dimensional representation of four biomolecular complexes tested in this study: protein-RNA (1NYB), protein-small-molecule (5N8V), DNA-small-molecule (6JJ0), and protein-protein (2W84). **b.** Mean binding-energy estimates over 100 frames per complex, reported in kcal/mol. The IE and C2 entropy rows show GB-OBC2 binding free energy estimates, including the corresponding entropy contribution. QM/MMGBSA calculations were performed for the two small-molecule complexes, treating residues within 4 Å of the ligand with quantum mechanics. **c.** Effect of different treatments on binding energies: ALPB correction (left), linear versus non-linear PB (center), differences in energies including the IE and C2 entropies (right). Created in BioRender. Valdes, M. (2026) https://BioRender.com/tehepwd

We illustrate gmx_MMPBSA’s capability to handle multiple calculation options and a diverse set of biomolecular complexes, including protein-RNA, protein-small molecule, DNA-small molecule, and protein-protein complexes. Calculations over the same 100 frames for each complex demonstrate how solvation treatments, considering explicit receptor waters, and including entropy estimates affect the calculated binding energies. The following sections describe these additions and improvements in detail.

#### Solvation models

The GB, PB, and 3D-RISM options available in the initial release have been extended with additional solvation treatments and calculation controls. Version 1.5.0 introduced support for solving the nonlinear PB equation through the PBSA solver in sander. Version 1.5.2 added the analytical linearized Poisson-Boltzmann (ALPB) approximation,^17^ which accounts for the solute-solvent dielectric contrast within an analytical electrostatic treatment. Support for the GBNSR6 model was introduced in version 1.6.0. This model computes effective Born radii numerically through an R6 integral over the solute molecular surface and is available in gmx_MMPBSA for energy calculations as well as per-residue and pairwise decomposition.^15^ The 3D-RISM workflow was expanded to expose additional AmberTools controls and support PC+ corrections.^18^ Since version 1.5.2, 3D-RISM calculations have been performed through sander, with precalculated solvent susceptibility (.xvv) files provided to facilitate their use.

We demonstrate the use of the new solvation models and methods to calculate effective binding energies (no entropy correction) across four classes of biomolecular complexes (Figure 3). For example, adding the ALPB correction to GB-OBC2 increased the calculated binding energy of the protein-RNA complex from −66.33 to −48.96 kcal/mol, whereas the protein-small molecule binding energy remained nearly unchanged (−55.54 vs. −55.65 kcal/mol). Similarly, linear and nonlinear PB yielded substantially different energies for the protein-RNA complex (−86.59 vs. −53.38 kcal/mol), with a smaller difference for the protein-small molecule complex (−53.42 vs. −50.35 kcal/mol). GBNSR6 energies were similar to those of the GB-OBC2 model, except for the protein-RNA complex, which showed an absolute difference of 6.37 kcal/mol. The larger shifts in the binding energy for the protein-RNA complex illustrate how sensitivity to these solvation treatments differs across systems.

#### Entropy estimation

Version 1.5.0 introduced the second-order cumulant (C2) entropy approximation,^19^ complementing the existing normal-mode and interaction entropy (IE)^20^ approaches. C2 entropy estimates the entropic contribution to binding free energy as:

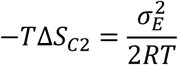

where 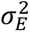 is the variance of the molecular-mechanics interaction energy over the selected frames, R is the molar gas constant (0.001987 kcal·mol^-1^·K^-1^), and T is the absolute temperature. Both IE and C2 report the standard deviation and standard error across deterministic, non-overlapping blocks, together with the block size and number of blocks. These diagnostics allow users to assess variability within the sampled trajectory.

Across the four biomolecular complexes included in this study, we showed how the choice of entropy estimator affects the resulting GB-OBC2 binding free energy estimates (Figure 3). For the protein-small molecule and DNA-small molecule complexes, the IE- and C2-corrected estimates differed by less than 1 kcal/mol. In contrast, the protein-RNA complex yielded −59.47 kcal/mol with IE and −50.88 kcal/mol with C2, a difference of 8.59 kcal/mol. The larger difference for the protein-RNA complex revealed entropy estimation as a particularly influential calculation choice in this example.

#### QM/MMGBSA

The QM/MMGBSA workflow has been extended to simplify quantum-region definition and reduce charge-assignment errors. Version 1.5.0 introduced automatic determination of the complex, receptor, and ligand QM-region charges, together with more flexible residue selection. For binding energy calculations, these charges must satisfy:

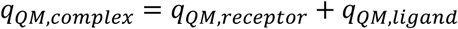

The selected quantum region is treated using the specified quantum method, while the rest of the system is described by molecular mechanics. Version 1.7.0 adopted PM6-DH+^21^ as the default method and checks the AmberTools calculation output for self-consistent field (SCF) convergence. When SCF convergence fails, the calculation stops immediately, and diagnostic information is provided to the user. This prevents the failed calculation from contributing to the reported binding energy estimate and avoids wasting calculation time.

We applied the updated QM/MMGBSA workflow to the protein-small-molecule and DNA-small-molecule complexes using PM6-DH+ and a QM region selected from residues within 4 Å of the ligand (Figure 3). Over the same 100 frames, QM/MMGBSA calculations yielded binding energy estimates of −11.76 and −21.08 kcal/mol, respectively, compared with −55.54 and −57.71 kcal/mol using GB-OBC2 calculations. In both examples, the QM/MMGBSA estimates were closer to the reported experimental binding free energies of −9.12 (K*i*=207nM)^22^ and −10.16 kcal/mol (K*d*=36nM),^23^ respectively; although a substantial discrepancy remained for the DNA-small molecule complex.

#### Membrane calculations

Membrane workflows use PB models with uniform or heterogeneous slab-like dielectric profiles.^24–27^ Membrane placement in MMPBSA calculations has been a recurrent issue among gmx_MMPBSA users, since it requires manual inspection of the system to determine the center (*mctrdz* parameter) and the approximate thickness (*mthick* parameter) of the membrane. Version 1.7.0 added automatic estimation of these parameters from selected lipid atoms (P atoms by default) in the complex trajectory. The membrane center is calculated from the mean z coordinate of these atoms over the analyzed frames, and the thickness is calculated from the difference between the mean z coordinates of atoms on either side of the center. The calculated parameters are stored in a CSV file, and a diagnostic plot is exported for user inspection (Table S1, Figure S1). This reduces manual parameter specification and allows users to verify the stability of the membrane placement.

#### Explicit waters

Water molecules are commonly removed when preparing the systems for end-state binding-energy calculations. However, previous studies have shown that retaining a hydration shell around the ligand or keeping interfacial waters can improve agreement with experimental binding affinities in some systems.^10,28–32^ Version 1.7.0 introduced an explicit receptor-water workflow that retains a specified number of waters near a selected reference region and includes them in both the complex and receptor calculations. By default, the reference region comprises interface residues identified from the change in solvent-accessible surface area between the isolated components and the complex. Alternatively, users can define the region through a receptor-ligand distance cutoff or an explicit residue selection. This option is available for single-trajectory GB, GBNSR6, PB, 3D-RISM, normal-mode, and QM/MMGBSA workflows. Water selection is performed dynamically, allowing the identities of the retained water molecules to change along the trajectory.

The binding energies obtained from GB-OBC2 calculations considering the 10 closest receptor waters around the interface differed from the standard GB-OBC2 results across all four complexes (Figure 3). For example, the protein-protein estimate changed from −53.99 to −29.14 kcal/mol, whereas the protein-RNA estimate changed from −66.33 to −62.00 kcal/mol. These calculations illustrate how the inclusion of selected receptor waters can affect binding energy estimates depending on the molecular system.

#### Computational mutagenesis

Alanine/glycine scanning is used to estimate the energetic effect of residue substitutions by comparing the energy and entropy terms in the native and alanine/glycine mutant systems.^33–37^ Earlier gmx_MMPBSA releases supported only single-residue substitutions. The latest versions (1.7.0 and 1.7.1) extended this functionality to multiple mutations, allowing several residues to be replaced simultaneously in one calculation. All selected residues must belong to the same molecular component and be replaced by the same residue type, either alanine or glycine (https://valdes-tresanco-ms.github.io/gmx_MMPBSA/1.7.1/examples/Alanine_scanning).

### Graphical and programmatic analysis

#### Analyzer redesign and improved performance

The redesign of gmx_MMPBSA_ana in the 1.5.x series improved the loading, organization, and display of calculation results. Parallel loading, optimized parsing, and optional disk-backed storage were implemented in gmx_MMPBSA_ana to handle longer trajectories and multiple systems more efficiently. In the new implementation, loading a file containing 200,000 energy values took 17 s, compared with 960 s previously. Loading time remained approximately unchanged when the number of systems increased to four, whereas the previous implementation required 4,515 s. For results containing both energies and pairwise decomposition, loading one and four systems took 9 and 14 s, respectively, compared with 192 and 960 s previously (https://valdes-tresanco-ms.github.io/gmx_MMPBSA/1.7.0/analyzer). These improvements make large result datasets and comparisons across multiple systems more accessible for interactive analysis.

#### API modernization and Jupyter notebooks

The Python API has been modernized to provide a consistent interface for graphical and programmatic analysis. Version 1.7.0 provided a loading function that returns a result object with dedicated methods for accessing energy, entropy, decomposition, and calculation metadata. Completed calculations can be loaded from a `_GMXMMPBSA_info` record and its associated files or from a portable, compact `.mmxsa` file:

~~~
from GMXMMPBSA import API
result = API.load(“COMPACT_MMXSA_RESULTS.mmxsa”)
energy = result.get_energy()
decomposition = result.get_decomp_energy()
~~~

The API returns energy, entropy, and decomposition data in a Pandas dataframe, as well as input settings, file information, and metadata. Using a compact `.mmxsa` file simplifies the transfer of completed results, whereas optional disk-backed storage reduces memory requirements when loading larger datasets.

Following the modernization of the API, we included two Jupyter notebooks (https://valdes-tresanco-ms.github.io/gmx_MMPBSA/1.7.0/examples/#jupyter-notebooks) to illustrate the use of the gmx_MMPBSA suite in local and cloud environments. The local notebook illustrates how to load the results through the API, extract energy and decomposition data, and generate plots that users can adapt to their own systems. The Google Colab notebook shows how to install the software and run gmx_MMPBSA examples in a hosted session. These notebooks document the commands and data-processing steps used in each analysis and provide a starting point for users interested in analyzing results programmatically.

## Availability

gmx_MMPBSA is distributed under the GNU General Public License v3.0. The source code, tagged releases, example datasets, and Jupyter notebooks are available on the GitHub repository https://github.com/Valdes-Tresanco-MS/gmx_MMPBSA. Online documentation for the latest version 1.7.1 at https://valdes-tresanco-ms.github.io/gmx_MMPBSA/1.7.1 provides installation instructions, tutorials, and examples covering supported molecular systems and calculation workflows.

## Conclusions and future directions

The gmx_MMPBSA suite offers an end-to-end workflow for performing end-state binding free energy calculations. Here, we report on the addition of new calculation protocols, native AMBER input support, and better analysis capabilities to the gmx_MMPBSA framework. The expanded suite retains compatibility with a broad range of GROMACS and AmberTools versions, which increases gmx_MMPBSA applicability. Using a diverse set of biomolecular complexes, we leveraged the new gmx_MMPBSA capabilities to illustrate how the solvation treatment, entropy estimation, and inclusion of explicit receptor waters affect the binding energy.

Future development will extend native input support to simulation engines such as OpenMM and NAMD, incorporate additional implicit-solvent treatments, and introduce complementary binding affinity approaches, including Linear Interaction Energy (LIE).

## Associated Content

### Supporting Information

Details on system preparation, MD simulation procedures, and energy calculations are described. Diagnostic data for automatic membrane placement (Table S1, Figure S1). Data and programming scripts used to reproduce Figures 2 and 3 are available via figshare: https://doi.org/10.6084/m9.figshare.34518393

## Acknowledgments

Special thanks to the JetBrains company (https://www.jetbrains.com) for granting a free, open-source license to use their software.

## Abbreviations

MM/PB(GB)SA: molecular mechanics/Poisson-Boltzmann (generalized Born) Surface Area
3D-RISM: 3D-Reference Interaction Site Model
MD: molecular dynamics
IE: interaction entropy
API: application programming interface
ALPB: analytical linearized Poisson-Boltzmann
C2: second-order cumulant
GB-OBC2: generalized Born model using the second Onufriev-Bashford-Case parameterization
PC+: modified pressure correction
QM/MMGBSA: quantum mechanics/molecular mechanics generalized Born surface area
SCF: self-consistent field

