## Supplementary Information for "Expanding gmx_MMPBSA capabilities for end-state binding free energy calculations with GROMACS and AMBER"

### System preparation and MD simulations

Systems were prepared using default options in CHARMM-GUI.^1^ Briefly, the AmberFF19SB force field^2^ was used for proteins, GAFF2 for small ligands,^3^ OL3 for RNA, and OL15 for DNA.^4^ The system was neutralized with 0.15 M NaCl and solvated with explicit water molecules, which were modeled by the OPC parameter set,^5^ in a rectangular box. The distance between any atom in a solute and the periodic box's edge was set to 10 Å. The system was minimized first with all heavy atoms restrained with 10 kcal/mol·Å^2^ to equilibrate the solvent. Production runs were performed in an NPT ensemble for 2 ns with position restraints. MD simulations were performed using the GROMACS (version 2022.3) software package.

### End-point binding free energy calculations

End-state binding free-energy estimates were calculated for the SARS-CoV-2 spike receptor-binding domain complexed with nanobody Nb20 (7JVB), the protein-RNA (1NYB), protein-protein (2W84), protein-small-molecule (5N8V), and DNA-small-molecule (6JJ0) complexes using gmx_MMPBSA. Calculations used the supplied GROMACS topologies, index selections, and solvated trajectories. Receptor and ligand groups excluded bulk solvent and ions. A single-trajectory protocol was used where the receptor and ligand coordinates were extracted from each complex snapshot, and binding-energy terms were calculated as the complex value minus the corresponding receptor and ligand values. Energies were summarized across the 100 frames as means with standard deviations and standard errors.

The implicit-solvent calculations for the SARS-CoV-2 spike receptor-binding domain complexed with nanobody Nb20 in gmx_MMPBSA, amber_MMPBSA, and MMPBSA.py used the GB-Neck2 (igb=8) model with *mbondii3* radii.^6^

The calculations for the rest of the complexes included GB-OBC2 (igb=5),^7^ GB-OBC2 with the ALPB correction (alpb=1),^8^ GBNSR6,^9^ and linear and nonlinear Poisson-Boltzmann models. All inputs used radii set *mbondi2* and an ionic strength of 0.15 M. PB, GB-OBC2, and ALPB used internal and external dielectric constants of 5.0 and 80.0, respectively. The Interaction Entropy (IE) and second-order cumulant (C2) entropy estimates were calculated separately from the GB-OBC2 energy calculations using the same GB parameters and 298.15 K temperature.

An additional GB-OBC2 calculation for each complex retained the 10 closest OPC waters as receptor waters, using the “within 4” interface selection. These calculations used an internal dielectric of 5.0. OPC extra-point sites were stripped; therefore, these results represent controlled relative comparisons rather than native OPC electrostatics.

For 5N8V and 6JJ0, QM/MMGBSA calculations were also performed with PM6-DH+, treating residues within 4 Å of the interface as the QM region and using GB solvation. Each run analyzed the same 100 frames as the other methods.

Table S1. Average z-value (center) and thickness across the selected frames. The data correspond to the protein-membrane example included in the gmx_MMPBSA documentation (<https://valdes-tresanco-ms.github.io/gmx_MMPBSA/dev/examples/Protein_membrane>)

| # selected_atoms=P | |  |  |  |  |
| --- | --- | --- | --- | --- | --- |
| # center_setting=automatic | | |  |  |  |
| # thickness_setting=automatic | | |  |  |  |
| # resolved_center_Å=62.315825 | | |  |  |  |
| # resolved_thickness_Å=37.298350 | | | |  |  |
| **frame** | **n_atoms** | **n_above** | **n_below** | **mean_z_Å** | **thickness_Å** |
| 1 | 200 | 100 | 100 | 62.26095 | 37.3799 |
| 2 | 200 | 100 | 100 | 62.273 | 37.3544 |
| 3 | 200 | 100 | 100 | 62.2199 | 37.2044 |
| 4 | 200 | 100 | 100 | 62.50945 | 37.2547 |

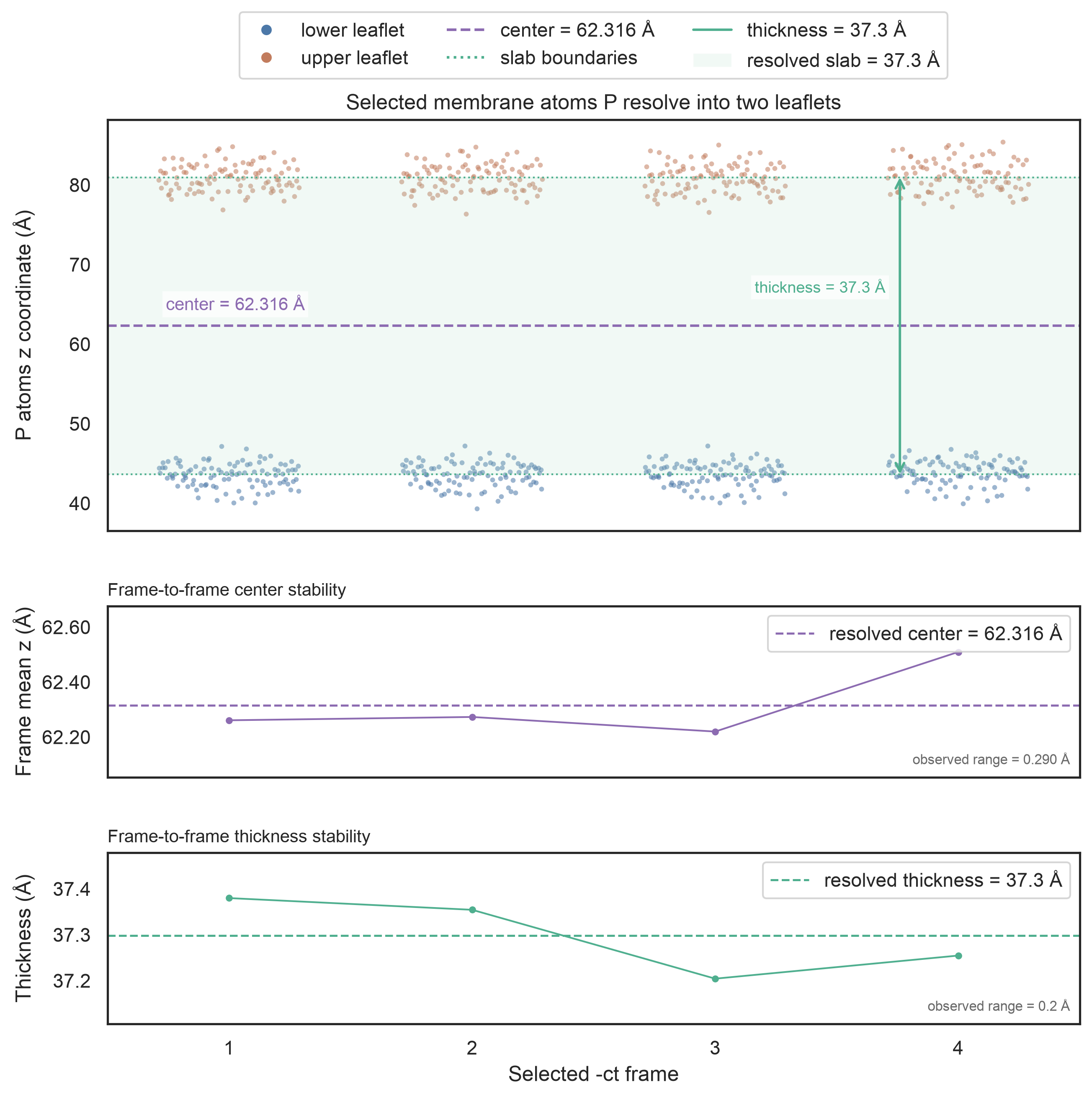

Figure S1. Diagnostic plots for automatic detection of *mctrdz* and *mthick* parameters. Top: variation in the z-coordinates of the selected membrane atoms (“P” atoms by default) in both leaflets is used to calculate the center (*mctrdz*) and thickness (*mthick*) parameters across the selected frames. Center and bottom: The average z-value (center) and thickness across the selected frames are used to calculate the center (*mctrdz*) and thickness (*mthick*) parameters. The data correspond to the protein-membrane example included in the gmx_MMPBSA documentation (<https://valdes-tresanco-ms.github.io/gmx_MMPBSA/dev/examples/Protein_membrane>)
